# Elastic fibers define embryonic tissue stiffness to enable buckling morphogenesis of the small intestine

**DOI:** 10.1101/2023.07.18.549562

**Authors:** Elise A. Loffet, John F. Durel, Richard Kam, Hyunjee Lim, Nandan L. Nerurkar

## Abstract

During embryonic development, tissues must possess precise material properties to ensure that cell-generated forces give rise to the stereotyped morphologies of developing organs. However, the question of how material properties are established and regulated during development remains understudied. Here, we aim to address these broader questions through the study of intestinal looping, a process by which the initially straight intestinal tube buckles into loops, permitting ordered packing within the body cavity. Looping results from elongation of the tube against the constraint of an attached tissue, the dorsal mesentery, which is elastically stretched by the elongating tube to nearly triple its length. This elastic energy storage allows the mesentery to provide stable compressive forces that ultimately buckle the tube into loops. Beginning with a transcriptomic analysis of the mesentery, we identified widespread upregulation of extracellular matrix related genes during looping, including genes related to elastic fiber deposition. Combining molecular and mechanical analyses, we conclude that elastin confers tensile stiffness to the mesentery, enabling its mechanical role in organizing the developing small intestine. These results shed light on the role of elastin as a driver of morphogenesis that extends beyond its more established role in resisting cyclic deformation in adult tissues.

## INTRODUCTION

The function of organs in the body is inextricably linked to their anatomical form. During embryonic development, these complex forms arise stereotypically from a remarkably simple starting point: a seemingly disorganized ball of cells. There has been great progress in developmental biology over the past few decades in understanding this process of organogenesis from a mechanical standpoint [1,2], focused on how forces are generated, and how they bend, twist, extend and shorten developing tissues across multiple length and time scales into their functional adult forms. Increasingly, the focus has shifted from purely mechanical views on morphogenesis to those that aim to link tissue-scale mechanics to the underlying biology that defines them [3]. The greatest advances in this vein have focused on understanding how cells in an embryo generate forces [4] and transmit them across tissues [5]. Importantly, for such forces to induce stereotyped tissue deformations, the material properties of embryonic tissue must be precisely and dynamically regulated throughout development. However, what determines the material properties of embryonic tissues has received relatively little attention until recently [6–8]. In the present study, we asked what controls the material properties of the dorsal mesentery, a tissue in the developing embryo that drives packing of the lengthy small intestine into the confines of the body cavity via buckling instability.

Within the adult digestive system, the small intestine is responsible for the greater part of nutrient absorption. This is achieved through maximizing the absorptive surface area of the gut, which in turn is accomplished through two morphological adaptations: small finger-like projections called villi on the luminal surface of the intestine, and gut looping, a process by which the intestine is organized into loops that allow its length to far exceed the linear dimensions of the body cavity. Errors in the formation of intestinal loops during embryonic development result in catastrophic birth defects. These include midgut volvulus, in which the intestine becomes entangled with itself, obstructing the bowel and strangulating the vasculature [9]; in its most extreme presentations, babies are born vomiting bile due to an inability to pass contents through their intestine. A second set of congenital disorders, including omphalocele and gastroschisis, involve a failure of the intestine to be properly internalized within the body cavity prior to birth, resulting in babies born with their intestines still outside the body [10,11]. The biological basis for these devastating gastrointestinal defects is not well understood, owing in part to a gap in our understanding of the molecular and genetic controls on normal looping morphogenesis of the small intestine. On the other hand, the mechanics of gut looping have been well described [12]. The initially straight intestinal tube elongates against the constraint of the dorsal mesentery, a thin membranous tissue attaching the tube to the dorsal body wall.

While both tissues elongate, the tube elongates much more rapidly than the mesentery, such that the mesentery is passively stretched by the tube, and in turn the tube is compressed by the mesentery [12]. This compression ultimately leads to a mechanical instability in the intestinal tube, buckling the tube into loops. The resulting wavelength and curvature of intestinal loops is highly reproducible within a given species, and can be accurately predicted using simple scaling laws based on geometry, stiffness, and growth rates of the tube and mesentery [12]. A similar number of loops in the intestine of all individuals of a given species suggests that material properties of the mesentery must be tightly regulated during organogenesis of the small intestine. Too soft a mesentery would fail to resist the elongating intestinal tube enough to reach a critical buckling load, while too stiff a mesentery would lead to excessive and abnormal looping consistent with birth defects such as midgut volvulus. Therefore, the stiffness of the mesentery must be precisely regulated during embryonic development to ensure the stereotyped morphogenesis of intestinal loops. While we have recently identified signals that regulate differential growth rates between the tube and mesentery [13], the biological determinants of tissue stiffness in the mesentery have not previously been identified. Indeed, while buckling has increasingly become recognized as a core physical mechanism shaping many organs in development [14,15], in most cases the determinants of tissue stiffness that effectively translate differential growth between two tissues into buckling forces have been understudied.

More broadly, there is a gap in our understanding of how tissue-scale mechanics are integrated with underlying cell and molecular effectors of morphogenesis [16]. While traditionally the focus of bridging this gap has been on understanding force generation [4,17,18], force transmission through cell-cell adhesion [19,20], or cell migration [21], it is only recently that extracellular matrix (ECM) has been recognized as an active participant in morphogenesis. For example, the ECM has been implicated in directing anisotropic growth of the Drosophila egg chamber [22] and early mammalian embryo [23], the initial chiral rotation of the primitive intestinal tube [24–26], and branching morphogenesis of mammary [27] and airway epithelia [28].

Material properties of the extracellular matrix (ECM) are mainly determined by the organization and enrichment of fibrous proteins [29], such as collagens, fibronectin and elastin. Elastin is the main element of elastic fibers, which together with scaffolding fibrillin proteins [30], form a highly-crosslinked network that functions to diffuse stresses during large deformations [31], enabling stretch and recoil in many soft tissues [32], including most notably, the arterial walls [33]. Previous studies into the mechanical role of elastin in various tissues have shown that depleting elastin can result in lower tensile modulus [34] as well as a reduced capacity to withstand load, resist fatigue loading, and recover after loading [35,36]. The biomechanical role of elastin has largely been studied in tissues that experience cyclic mechanical loading as part of their physiologic function, such as cardiovascular tissue [33], the lungs [37], and connective tissues [38]. However, the mechanical role of elastin as a driver of tissue shape during embryonic development has not been appreciated.

In a clinical setting, known genetic defects in the elastin gene are for the most part linked to vascular diseases [39]. Mutations in the *ELN* gene [40] as well as other genes involved in elastic fiber formation [41] have also been associated with cutis laxa, a disease in which patients have abnormally loose skin. Other syndromes that are characterized with similarly loose connective tissues such as Marfan syndrome or Ehler-Danlos syndrome can also manifest with elastic fiber defects [42,43]. Case reports support a link between these syndromes and defects in the intestinal system such as intestinal volvulus, visceral ptosis or mobile caecum [44–46]. This suggests there could be a role for elastic fibers in the development and homeostasis of the gastrointestinal system, but such a link has not been tested directly.

During gut looping, the intestinal tube passively stretches the dorsal mesentery elastically to over 100% strain (reversibly stretching to double its unloaded length), and storage of this elastic energy within the mesentery is necessary to buckle the tube into compact loops. However, a role for elastin in looping morphogenesis of the small intestine has not been previously investigated. In the present study, we first broadly examined transcriptome wide changes in the mesentery during gut looping of the developing chick embryo, identifying upregulation of elastic fiber-related genes. Through subsequent functional studies, we observed that elastin plays a critical role in establishing the material properties of the dorsal mesentery, the central driver of buckling morphogenesis in the small intestine.

## RESULTS

### 1- Transcriptomic changes of the mesentery are dominated by upregulation of ECM genes during looping

Mechanical properties of the dorsal mesentery are central to buckling of the small intestine, but this tissue has been understudied from the biological standpoint - both during development and in adulthood [47]. Hence, in the absence of a strong contextual understanding of mesentery biology, we opted to perform a transcriptomic analysis of the tissue throughout the stages of looping, on the grounds that genes related to regulation of mesentery growth and stiffness are likely differentially regulated before, during, and at the end of looping. We performed bulk RNA-Seq analysis on mesentery tissue collected from chick embryos at embryonic day 8 (E8) when looping begins, E12 when looping is underway, and E16 when looping is complete (Figure 1a, b). Principal Component Analysis (PCA) revealed a clear separation of transcriptomes by developmental stage along the first principal component, which accounted for approximately 52% of the variance (Figure 1c). Although the second principal component captured separation between the mid-looping E12 transcriptome and that of E8 and E16, this appeared to be primarily due to transient regulation of genes associated with development of the peripheral nervous system (Supplemental Figure 1). We next performed differential expression analysis to extract differentially expressed genes through time (thus correlating with progressive looping of the intestine). Through Gene Ontology analysis of these results, we found that genes upregulated through time were strongly linked to ontologies related to ECM gene expression, including for example, “collagen-containing extracellular matrix”, “extracellular matrix” or “extracellular matrix organization” between E8 and E12 (Figure 2a), and “extracellular space”, and “extracellular region” genes between E12 and E16. At E16 when looping concludes, several collagen genes were upregulated, including *Col1a1*, *Col1a2*, *Col4a1*, *Col8a1*, *Col6a1* (Figure 2b). Interestingly, several genes encoding proteins involved in elastic fiber formation were also upregulated, including genes encoding the core protein of elastic fibers, *Eln* (tropo-elastin); an accessory protein involved in elastic fiber assembly, *Fbln5* (fibulin-5); and an enzyme responsible for crosslinking elastin and collagen fibers, *Loxl2* (lysyl-oxidase) (Figure 2c). Elastic fibers are attributed with conferring high extensibility to load-bearing tissues [48]. Because the mesentery drives buckling of the intestinal tube by accommodating in vivo strains as high as 150% elastically, we focused next on investigating the role of elastin in prescribing mechanical properties of the mesentery that produce stereotyped buckling of the intestine by resisting its elongation [12].

**Figure 1.**
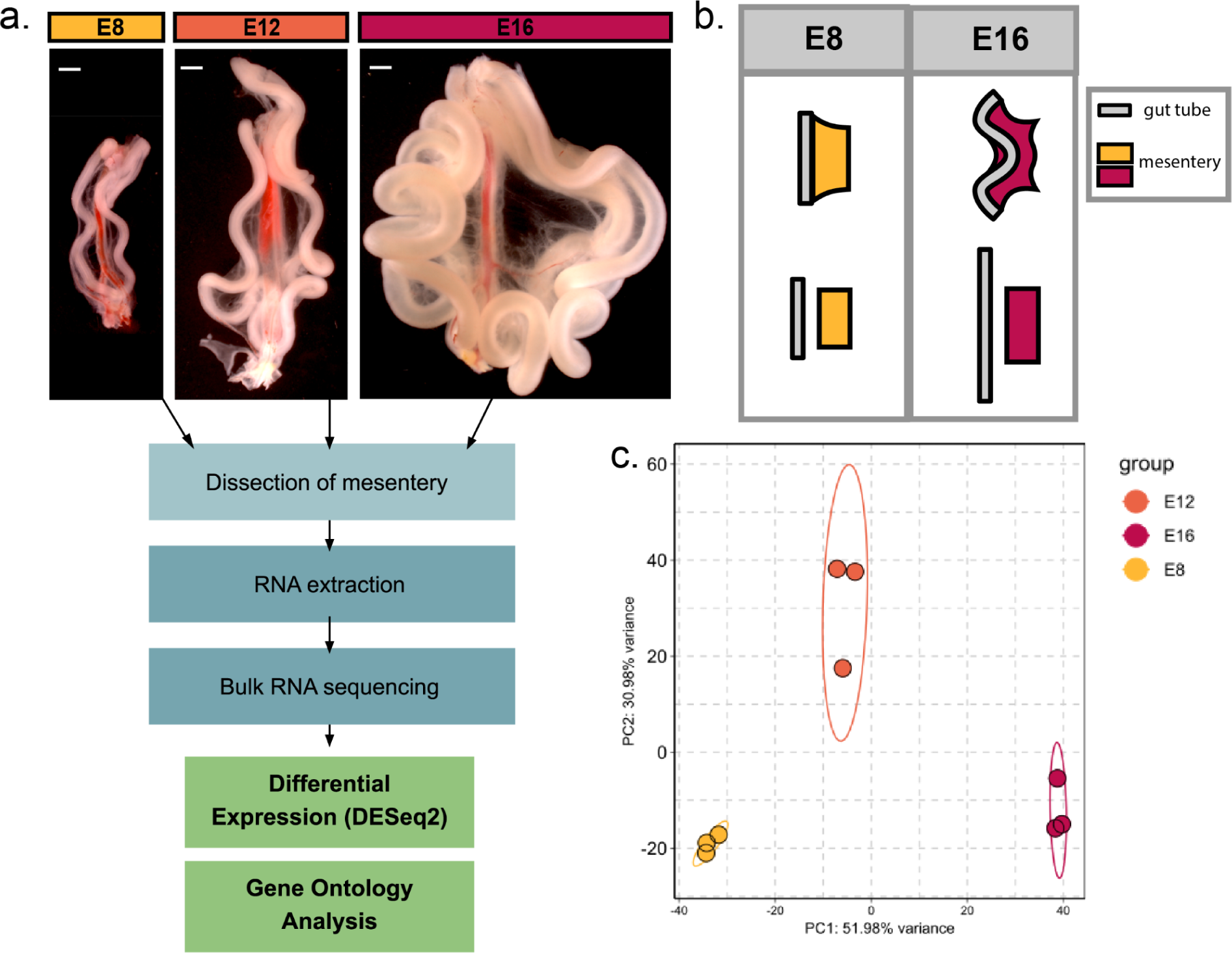
Overview of bulk RNA sequencing analysis of the dorsal mesentery during buckling morphogenesis of intestinal loops. **a.** Intestines from chick embryos at E8, E12 and E16, reflecting the beginning, middle, and end, respectively, of buckling morphogenesis of intestinal loops (scale bar = 1mm); below, a schematic overview of the bulk RNAseq experiment. **b.** Graphic representation of buckling of the intestinal tube driven by differential growth between the tube and the mesentery; separated lengths of the tube and mesentery are shown below the intact configurations to reflect accentuation of differential elongation rates over time. This growth differential drives buckling of the intestine into loops. **c.** Principal Component Analysis on 1000 genes of bulk RNAseq data of mesentery tissues at E8, E12 and E16.

**Figure 2.**
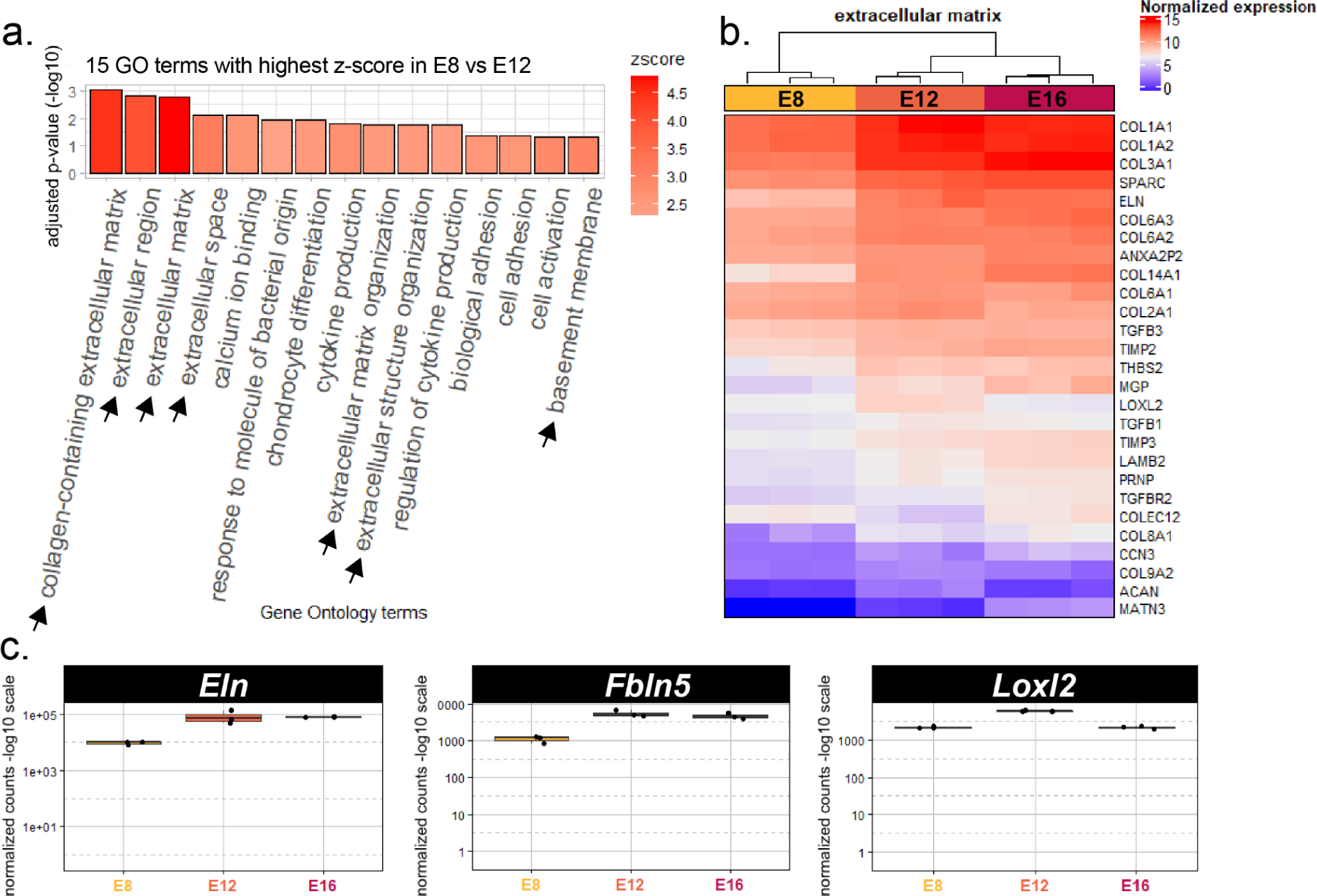
Bulk RNAseq reveals broad transcriptome changes related to extracellular matrix during looping. **a.** Bar plot of 15 Gene Ontologies with highest E12-associated z-score for E8 vs E12 differential gene expression analysis. Z-score is calculated based on how many genes in the gene ontology are up-regulated at E12 vs E8. Arrows indicate ontologies linked to the extracellular matrix. **b.** Heatmap of normalized expression of genes in “extracellular matrix” gene ontology that are differentially expressed between developmental stages. **c.** Box plots of normalized read counts for *Eln*, *Fbln5*, *Loxl2*.

### 2- Elastic fiber formation and mechanical properties of the dorsal mesentery

Building on the transcriptomic data, we first examined the localization and relative abundance of elastin protein in the mesentery. Immunofluorescence staining for elastin showed a progressive increase in elastin protein in the mesentery from E8 to E16 (Figure 3a). Although elastin was observed within the intestinal tube surrounding blood vessels and bounding the circumferential smooth muscle layer, it was particularly enriched in the mesothelium, a thin epithelial layer that separates the intestinal tube from its surrounding coelomic cavity (Figure 3a). We next carried out a baseline characterization of mesentery mechanics under uniaxial tension, employing a custom micromechanical tester to examine how properties change from the onset of looping at E8 through completion of looping at E16. The mesentery has a nonlinear stress-strain behavior, with a low-stiffness toe region followed by a transition to a high-stiffness linear region (Figure 3b). Accordingly, mesentery mechanics were characterized by measurement of a toe-region modulus E_toe_, transition strain ε* denoting the strain at which the material transitions between soft and stiff response, and the linear region modulus E_lin_. We observed no changes in the toe-region modulus E_toe_ (Figure 3c) or transition strain ε* (Figure 3c) over the course of looping. However, E_lin_ increased over five-fold from the onset of looping at E8 to its completion at E16 (Figure 3d). To consider these mechanical changes in the context of the temporal progression of elastic fiber assembly, we next examined the expression of *Eln* by qPCR. *Eln* expression increased markedly from E8 to E16 (Figure 3e), correlating with E_lin_ (Figure 3f), but not E_toe_ or ε* (Supplemental Figure S2) over time. Together, these data suggest that elastic fibers may play a role in establishing the large-strain stiffness of the mesentery during looping. We next tested whether this is the case by studying the effects of elastin depletion on tissue mechanics.

**Figure 3.**
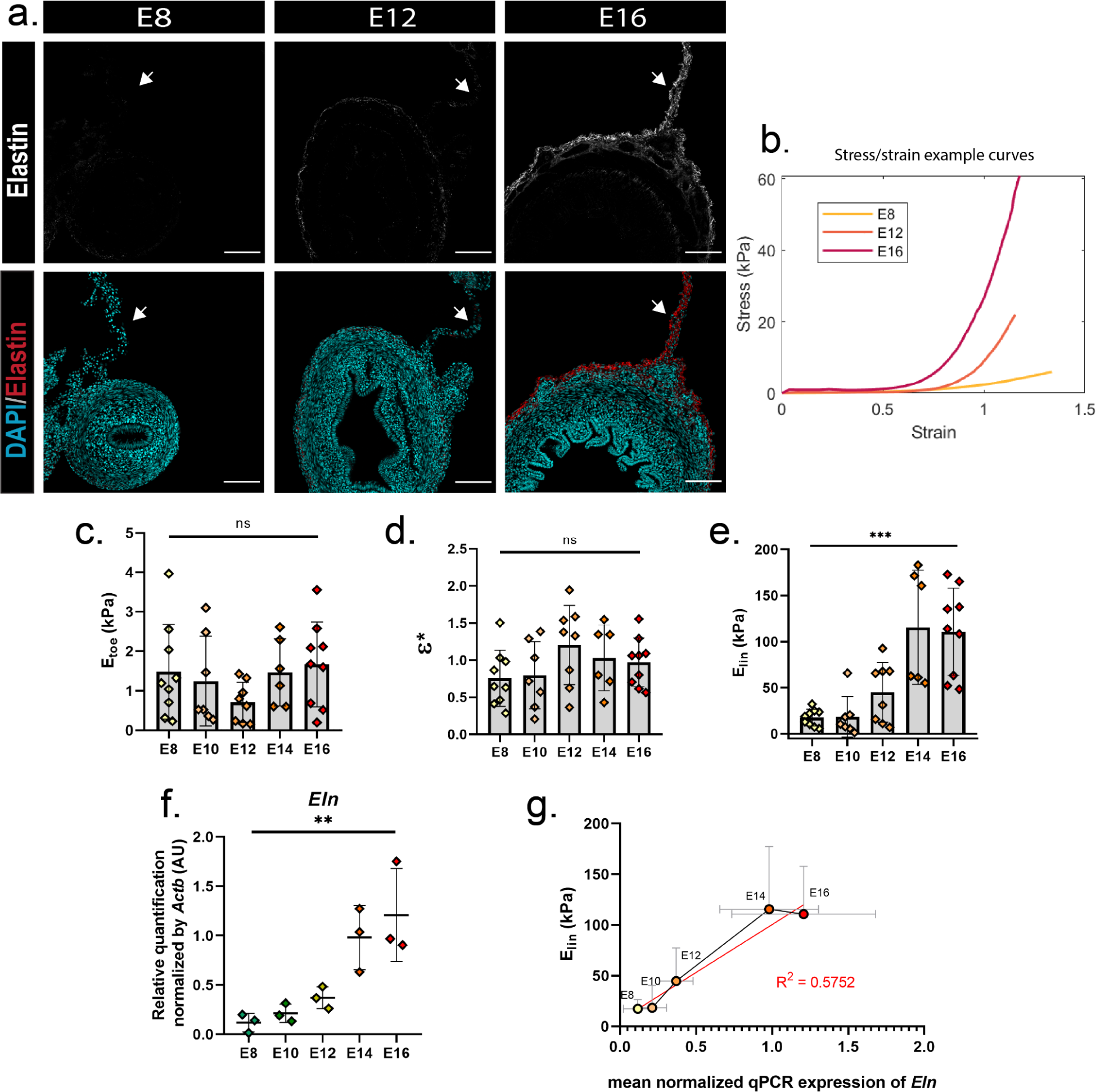
Elastin protein is enriched in the mesentery during looping and expression levels correlate with tissue stiffness. **a.** Immunofluorescence staining for elastin and cell nuclei (DAPI, blue) on E8, E12 and E16 sections (scale bar = 100 μm). White arrowhead indicates dorsal mesentery. **b.** Representative stress-strain curves from uniaxial tensile testing of mesenteries at E8, E12, and E16. **c-e.** Toe modulus (E_toe_, c), transition strain (ε*, d) and linear modulus (E_lin_, e) for mesenteries calculated from uniaxial tensile tests of mesentery between E8 and E16; for E_lin_, p < 0.0001 for E8 vs E16, p< 0.05 for all other comparisons except E8 vs E10 and E14 v E16). **f.** qPCR for *Eln* in the mesentery between E8 and E16 (p < 0.005 for E8 vs E16 and E10 vs E16, p < 0.05 for E8 vs E14, E10 vs E14, E12 vs E16). **g.** Correlation between expression levels of *Eln* and E_lin_ from E8 to E16 (R^2^ =0.5752). *ns = not significant (p>0.05); *p <0.05; **p <0.005; ***p <0.0005*

### 3- Selective removal of elastic fibers from the mesentery

Based on the correlation over developmental stage between mesentery stiffness and elastin protein and gene expression, we next asked whether elastic fibers play a functional role in establishing the mechanical properties of the mesentery, and by extension whether elastin regulates buckling morphogenesis of the small intestine. To test this, we relied on selective degradation of elastin using the proteolytic enzyme elastase [49], which has been widely utilized to study the role of elastin in adult tissues such as tendons [35], arterial wall [50] and intervertebral disc [34], among others. An initial dosage study revealed that incubation with 2 or 4 U/mL of elastase for 4 hours at 37°C depleted elastin in the mesentery at E16, when looping is complete (Figure 4a). Live-dead staining revealed that mesentery cells remained healthy through the course of treatments, with no notable increase in cell death compared to controls (Figure 4b, Supplemental Figure S3a). To reduce the potential for off-target effects on collagen, elastase treatment was performed in the presence of 0.1mg/mL of Soy Bean Trypsin Inhibitor (SBTI) [51]. Nonetheless, to verify that off-target effects on collagen were minimized, we examined collagen organization and abundance following elastase treatment. Second Harmonic Generation (SHG) imaging of the mesentery revealed that, while some subtle changes in collagen organization may be observed at 4 U/mL elastase, overall, collagen remained abundant within the mesentery when compared to controls (Figure 4c). This was further verified quantitatively by biochemical analyses of collagen content, which revealed no loss of collagen with elastase treatment (Supplemental Figure S3b). Together, these findings indicate that elastase treatment effectively removes elastin from the mesentery without compromising cell viability or collagenous ECM.

**Figure 4.**
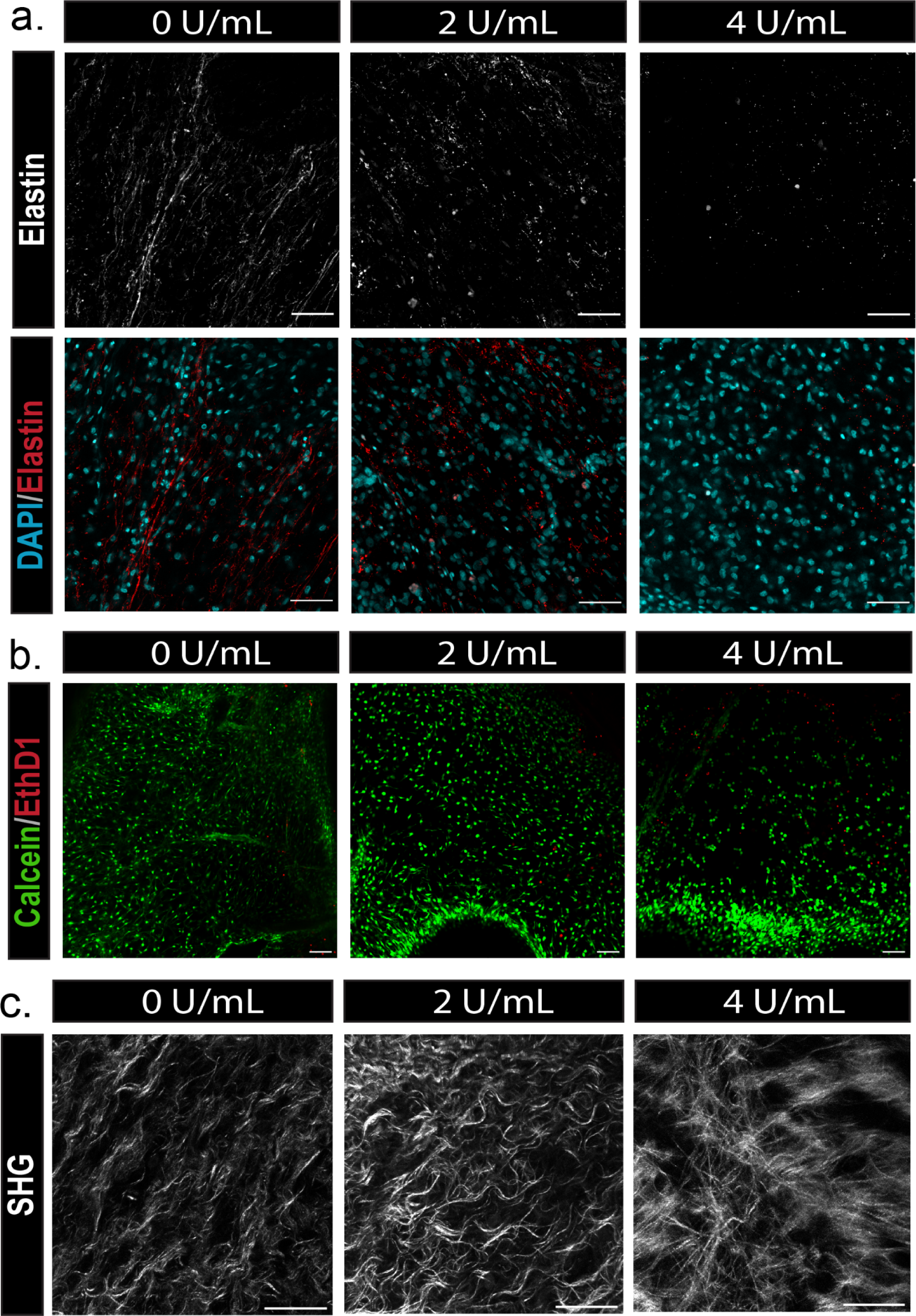
Elastase treatment degrades elastin while preserving cell viability and collagen organization in the E16 mesentery. **a.** Whole-mount immunofluorescence staining for elastin and DAPI in the mesentery following incubation of intestine and mesentery segments with 0, 2, or 4 U/mL elastase (scale bar = 50 μm) **b.** Live/dead staining to assess cell viability, where live cells stain positively for calcein (green) and dead cells stain for ethidium homodimer 1 (EthD1, scale bar = 100 μm) **c.** Second Harmonic Generation (SHG) imaging of collagen network in control and elastase-treated mesenteries (scale bar = 50 μm).

### 4- Elastin depletion reduces mesentery modulus

We next examined whether depletion of elastin from the mesentery directly influences tissue mechanics. Interestingly, elastase treatment reduced both the toe and linear region moduli (Figure 5a, c), but did not alter the transition strain ε* (Figure 5b). The effects on linear modulus E_lin_ were particularly dramatic, with a decrease of approximately 80% observed upon treatment with either 2 or 4 U/mL elastase. To gain further insight into the role of elastin in determining material properties of the mesentery, we quantified the effect of elastase treatment on the Poisson’s ratio. The mesentery displays a strongly nonlinear Poisson behavior, with Poisson’s ratio increasing with stretch (Figure 5d). No changes in Poisson’s ratio were observed upon treatment with elastase in either the toe or linear region (Figure 5d), indicating that nonlinear Poisson behavior is unchanged with elastin depletion.

**Figure 5.**
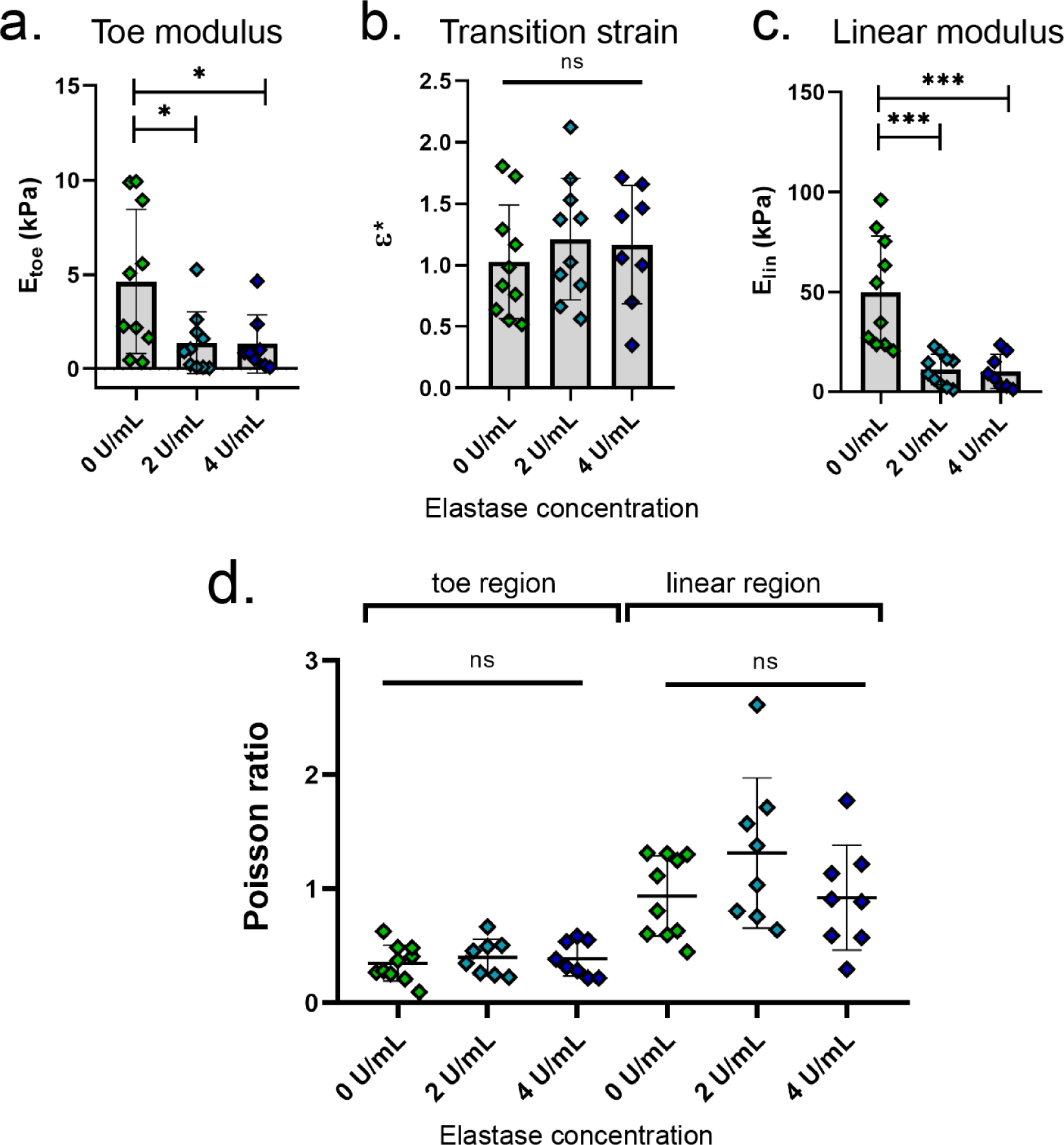
Elastin depletion reduces toe and linear moduli of the mesentery. **a-c.** Toe modulus (E_toe_, a), transition strain (ε*, b), and linear modulus (E_lin_, c) for control and elastase-treated mesenteries. **d.** Poisson’s ratio for control and elastase-treated mesenteries measured while stress/strain curve was in the toe region, and in the linear region. *n.s. = not significant (p>0.05); *p<0.05; **p<0.005; ***p<0.0005*.

Together, these findings suggest that elastin plays an important role in defining the mechanical properties of the embryonic mesentery. Specifically, these results are consistent with elastin functioning to enable elastic energy storage within the mesentery as it is stretched by the elongating intestinal tube to very high strains. We therefore investigated energy dissipation in treated and untreated mesentery (Supplemental Figure S4a), and observed that elastase-treated samples exhibited an increase in hysteresis loss (Supplemental Figure S4b). This indicates that elastin may indeed play a role in energy storage in the mesentery. This test also showed an increase in deformation of the samples after unloading (Supplemental Figure S4c), in line with the described role of elastin in recoil and resistance of tissues [32].

## DISCUSSION

The ability of the dorsal mesentery to accommodate large deformations while providing the appropriate resistance is a critical determinant of intestinal looping in the embryo. In this study, we investigated the molecular effectors of this remarkable material behavior that allows buckling morphogenesis to occur so precisely during normal intestinal development. Beginning with a transcriptome-wide analysis of the mesentery, we observed that looping is associated with large-scale changes in gene expression, and most notably that many genes encoding ECM components are enriched as looping proceeds and the mesentery stiffens. This included genes associated with elastic fibers that are known to confer resistance and recoil to a range of adult tissues. We found that elastin depletion from the mesentery dramatically reduced the tensile stiffness in both the toe and linear region. This suggests that the ability of the mesentery to resist and therefore buckle the elongating intestinal tube is strongly dependent on elastin, and that elastic fibers have a previously unappreciated role in organogenesis of the small intestine.

Traditionally, elastic fibers have been observed and studied in tissues that experience large deformations under cyclic loading, such as large arteries and the lungs, where they act to enable large repeated deformations with full subsequent recovery. It was therefore unexpected that elastin transcripts and protein would be enriched in the mesentery, which is not subject to cyclic loading (outside of minimal passive extension from peristalsis of the intestine). Instead, the embryonic mesentery is primarily deformed under pseudo-steady conditions with 150% strain being progressively applied by the elongating intestine over the course of 8 days of development *in vivo*, a strain rate on the order of approximately 10^-6^/s. This suggests that elastin may have a unique biomechanical role during development that diverges from its more conventional and well-studied role in the function of adult tissues. Specifically, when tissues buckle during development as a consequence of differential growth, it is essential that the constraining tissue is able to accommodate large deformations without dissipating stored energy. Our results support the idea that elastin does exactly this in the mesentery, enabling the storage of elastic energy that then leads to buckling and maintenance of stable loops in the intestine. This is likely a common requirement to many organs in the body that are shaped by buckling due to constrained growth, including the cerebral cortex [52], branching airways [53,54], and of particular note, the intestinal villi [55]. Villi form on the luminal surface of the intestine due to growth of the inner layers of the tube against the surrounding stiff constraint of oriented smooth muscle layers within the walls of the intestinal tube. Interestingly, we observed elastic fibers within these very smooth muscle layers in the present study (Figure 3a), suggesting that elastin could also be regulating stiffness in the muscle layer and therefore supporting the notion that elastin’s role in looping morphogenesis in the intestine likely reflects a broader role in the buckling morphogenesis of a range of tissues and organs during vertebrate development.

Elastin depletion in the mesentery resulted in a reduction of the linear modulus by as much as 80%, indicating that elastin is a key contributor to tissue mechanics. However, the precise mechanism by which it functions is not yet clear. Elastin is roughly two orders of magnitude softer than collagen [56]. It was therefore unexpected that elastase would cause such a dramatic reduction in tensile properties, given that the collagen itself was largely unaffected (Figure 4c and Supplemental Figure S3b). One possible explanation is that elastic fibers enable load transfer between collagen fibers, as has been proposed in other tissues [57]. In this case, disruption of elastic fibers may lead to inefficient loading of collagen fibers, favoring fiber sliding over stretching, to produce an overall softer material response throughout the large deformations observed in the mesentery. This is consistent with studies in the adult intervertebral disc, where a similar reduction in toe and linear region moduli upon elastase treatment were attributed to a disruption of the elastin-collagen interactions [34]. While additional studies in the cardiovascular system [58] support a role for elastin in regulating the linear/large strain modulus, in other contexts, such as tendon fascicles [35] and ligaments [36], elastin depletion has shown no effect on linear modulus. These varied reports further indicate that the biomechanical function of elastin is context specific, and may vary between different soft tissues, or between embryogenesis and adulthood. Finally, it is noteworthy that, while such large changes in toe and linear region moduli were observed with elastin depletion, that the transition strain and Poisson’s ratio of the mesentery did not change. This suggests that these material properties are not only separable, but likely under control of distinct structural or cellular controls.

Our interest in mesentery mechanics during development ultimately relates to intestinal packing: regulation of mesentery stiffness directly controls the elastic resistance experienced by the elongating intestine, in turn determining the buckled morphology of the tube. We therefore examined the effects of elastase treatment on morphology of the intact intestine at E16, when looping is complete. However, despite the marked reduction in mesentery modulus, no significant changes were observed in loop number or morphology (Supplemental Figure S5). Because elastase treatment cannot be targeted exclusively to the mesentery, this likely reflects the fact that elastin was also degraded from the tube: a softer intestinal tube will have a lower bending stiffness, buckling at lower loads. Indeed, scaling laws derived previously suggest that equivalent changes in the moduli of tube and mesentery would effectively cancel out, and have no influence on buckling wavelength and curvature [12]. From these scaling laws, however, one can estimate the effects of isolated changes in the mesentery if it were possible to limit elastase treatment to the mesentery alone. All else equal, an 80% reduction in the linear modulus of the mesentery would reduce the number of loops in the E16 intestine from 10 to approximately 4. This is well beyond the natural standard variation observed in wild type embryos, and illustrates how the magnitude of changes effected by elastin depletion is highly significant from the physiologic standpoint.

Elastin depletion not only reduced tissue stiffness, but also resulted in an increase in hysteresis loss during mechanical testing (Supplemental Figure S4b). As an important caveat to these experiments, however, large hysteresis was also observed in control groups, despite stretching samples to a lower magnitude than what is observed in vivo during looping. This suggests that our micromechanical testing setup (where preload and preconditioning steps are not possible and the tests are conducted in a PBS bath) may not be appropriate for measuring viscoelastic effects, or that sufficient time was not given for full tissue recoil. Nonetheless, a dramatic increase in hysteresis and permanent deformation upon elastin depletion is consistent with elastin contributing to energy storage of the mesentery during looping. Energy storage by elastin has been associated with both its intra- and inter-molecular structures. Elastin’s structure has a high degree of conformational disorder, giving it a high entropy at relaxed state and making elastin highly flexible. Then, under tension, the high degree of cross-linking between elastic fibers permits the distribution of stresses throughout the tissue [32]. However, it is worth noting that the role of elastin in energy storage also appears to be context specific, with some studies reporting increased hysteresis loss upon elastin depletion [35], while others reported no effect [36].

While the precise morphology of intestinal loops is highly conserved for a given species, a range of morphologies are observed across species, likely reflecting evolutionary selection pressures that must balance diet with body size and shape. It is remarkable that the resulting morphological diversity of intestinal loops across birds and mammals arise through variations in physical, geometric, and growth properties of the tube and mesentery, while maintaining nearly the same magnitude of tensile stress in the mesentery across species [12]. This suggests the potential for a conserved mechanical feedback mechanism among amniotes, whereby tissue stiffness is regulated by stretch to maintain a ‘homeostatic’ state of stress on the resident cells of the mesentery. Indeed, *Eln* expression levels in the mesentery are correlated with mesentery tension, as expression levels increase during the phase of most rapid differential growth between the tube and mesentery (Figure 3f). Further, in light of the present work, it would be reasonable for mechanical feedback to act at the level of a molecule such as elastin that is central to setting tissue stiffness. *In vitro* studies have implicated mechanical activation in the regulation of *Eln* expression levels and elastic fiber deposition [59]; however these studies have primarily focused on the effects of cyclic loading on vascular smooth muscle cells. Future studies will investigate the extent to which elastic fibers in the mesentery function within a mechanical feedback loop to balance stretch and stress to ensure stereotyped morphogenesis of intestinal loops.

The elaboration of complex morphologies during embryonic development necessitates precise control over the material properties of developing tissues. Here, we examined the basis of mechanical properties in the dorsal mesentery, a tissue responsible for buckling the intestinal tube into compact loops through conversion of differential growth into compressive forces on the tube. Specifically, from a transcriptome-level analysis of tissue dynamics, we identified upregulation of ECM-related genes during looping, including genes associated with the formation of elastic fibers. Through selective degradation, we identified a critical role for elastin in establishing the large-strain stiffness of the mesentery, providing new insights on the role of elastic fibers in morphogenesis of the small intestine. These studies highlight the importance of combining molecular and genetic approaches with biomechanics to understand the physical basis of organogenesis, and reveal an important role for elastin in organogenesis of the small intestine.

## METHODS

### 1. Embryos

Fertilized White Leghorn chicken (*Gallus gallus domesticus*) eggs were acquired from the University of Connecticut Poultry Farm. Eggs were incubated at 37°C and 60% humidity. At E8, 12, and 16, eggs were opened, and embryos sacrificed by decapitation, followed by microdissection in PBS to isolate the small intestine for subsequent analyses.

### 2. Bulk RNAseq analysis

Mesentery tissues were dissected from sacrificed embryos at E8, E12, and E16. In brief, the intestinal tube and superior mesenteric artery were removed from the mesentery. Mesenteries were collected and flash frozen on dry ice. All dissections were done in PBS + DEPC (Diethyl Pyrocarbonate) with sterile tools. Tissues from 3 embryos were pooled for each replicate (3 for each condition) in order to reach the minimum tissue mass required. Samples were shipped as whole frozen tissue to Genewiz for RNA isolation (Qiagen RNeasy Plus Universal mini kit), library preparation (NEBNext Ultra RNA Library Prep Kit for Illumina), and sequencing (Hiseq). Quality control of sequenced data was performed using FastQC v0.11.9. Residual adapter sequences were cleaned using (cutadapt v3.1 (trim_galore extension)). Genome alignment was performed using hisat2 v2.2.1 (with chicken reference genome GRCg6a, Ensembl 105). A read count matrix was obtained with htseq v0.13.5. Subsequent analyses were carried out using R, including differential expression analysis with package DESeq2 (test : LikelyHood Ratio Test, threshold for genes to be considered expressed differentially : adjusted p-value < 0.05, log2Fold Change < -1 or > 1), PCA plot and gene loadings with pcaExplorer, and Gene Ontologies obtained with clusterprofiler (adjusted p-value cutoff : 0.05, q-value cutoff : 0.05).

### 2. Fluorescence microscopy

Samples were subject to immunostaining as described previously [60]. Briefly, whole dissected intestines were fixed in 4% PFA in PBS overnight at 4°C, then incubated in a sucrose gradient up to 30% sucrose before embedding in O.C.T compound (Scigen Scientific) and freezing on an ethanol dry ice bath. 16 μm tissue slices were cut using a Leica CM3050 S Cryostat. Sections were washed (PBS + 0.1% Triton), permeabilized (PBS + 0.3% Triton), incubated in blocking buffer (PBS + 0.1% Triton, 5% goat serum, 1% BSA, 1% DMSO) at room temperature, incubated overnight at 4°C in blocking buffer with primary antibody (1:500 mouse anti-Elastin, Millipore Sigma MAB2503), washed, and incubated with secondary antibody (1:500 Goat anti-Mouse Cy3), Jackson Immunoresearch 111-165-003) in blocking buffer for 1 hour at RT. Samples were counterstained in DAPI for 20 min at room temperature, washed in PBS, mounted in Fluoromount and coverslipped. Whole mount staining was performed similarly, with secondary antibody applied overnight at 4°C with agitation. Laser scanning confocal imaging was conducted on an inverted LSM880 microscope (Zeiss). For second harmonic generation (SHG) imaging of collagen structural organization, samples were dissected free of the attached intestinal tube, and imaged in whole mount. SHG signal was generated via two-photon excitation (Chameleon Ultra2, Coherent) at 850 nm, and emission detected at 425 nm using a 40X water immersion lens. Presented images are a maximum intensity z-projection (Produced with FIJI) of a 7µm thick z-stack.

### 3. Enzymatic treatments

Whole intestines or 1mm wide segments consisting of mesentery and attached intestinal tube were washed in PBS after dissection, and incubated in a solution of 0.1 mg/ml soybean trypsin inhibitor (Sigma-Aldrich) in PBS with varying concentrations of elastase (Elastin Products Company) for 4 hours at 37°C under agitation. Samples were then washed in PBS for 10 minutes and either fixed in 4% PFA for immunostainings or used for further experiments.

### 4. Mechanical testing

Mechanical properties of the dorsal mesentery were measured in uniaxial tension using a custom micromechanical tester that sits atop the stage of a Zeiss Axiozoom fluorescent macroscope, as described previously [13,55,61]. Briefly, 1 mm wide strips of mesentery and gut tube were cut from the intact gut, and thickness was measured optically. Samples were then hooked to a tungsten cantilever attached to a linear actuator; the cantilever serves both as a force transducer (by monitoring deflection) as well as a means to apply stretch to the sample. Fluorescent vital dye was used to place fiducial markers within the mesentery. Tracking of marker displacements and cantilever deflection during tensile tests was automated via custom code, and used to construct stress-strain curves, from which the toe-region modulus E_toe_ and linear region modulus E_lin_ were determined by bi-linear curve fits, with the transition strain ε* quantified as the intersection of the pair of linear regressions. All image analyses were performed in MATLAB.

### 5. qPCR

Total RNA was extracted from samples using RNeasy mini kit (Qiagen) and the resulting RNAs were reverse transcribed using SuperScript III Reverse Transcriptase (Life Technologies). Real-time quantitative PCR (qPCR) was performed using a StepOne thermocycler with a Sybr green mix kit (Applied Biosystems). Primer sequences were chosen for *Eln (*forward: GGTGTCCCAGGTGTGGTG, reverse: GAAGCGGATACCTGCTCCA) and housekeeping gene *Actb* (forward: CTGTGCCCATCTATGAAGGCTA, reverse: ATTTCTCTCTCGGCTGTGGTG)[62]. The relative expression of *Eln* was measured by the 2-ΔΔCt method and normalized for the housekeeping gene *Actb*.

### 6. Viability testing

Live/dead straining was conducted according to manufacturer’s instructions (L3224 live/dead staining kit, Invitrogen). Tissues were washed in PBS for 10 min and incubated in 2 μM calcein and 4 μM EthD1. Stainings were then imaged with an LSM880 confocal microscope (Zeiss).

### 8. Hydroxyproline (OHP) assay

Collagen content was quantified following a well-established biochemical assay for hydroxyproline [63]. Briefly, samples were frozen at -80°C, and freeze dried in a Labconco Lyophilizer for 2 days. Dried samples were weighed and digested with a 0.5mg/mL Proteinase K solution overnight at 56°C. The Proteinase K digest was then hydrolysed with HCl at 110°C overnight and left to dry at 40°C for 4 days. Hydrolyzed product was resuspended and OHP assay was performed by incubating samples and standard solution with Chloramine T at RT for 20 min, followed by Dimethylaminobenzaldehyde at 60°C for 15 min. OD values were measured with a μQuant (Biotek), and standard curve (R^2^=0.999) was used to determine collagen concentration.

### 9. Statistics

Statistical analyses were performed using one-way ANOVA test, with α=0.05. Analyses and visualization were produced with GraphPad Prism 8.0.1.

## AUTHOR CONTRIBUTIONS

EAL and NLN designed the study, wrote and revised the manuscript; HL and JFD collected tissue samples for RNAseq samples; JFD wrote mechanical testing analyses code; EAL performed all other experiments and analyses, with assistance from JFD in mechanical testing and second harmonic generation imaging, and assistance from RK for immunostaining and mechanical testing.

## DATA AVAILABILITY

The bulk RNAseq have been deposited in the Gene Expression Omnibus (GEO) under accession number GSE237492. Other raw data such as microscopy images and analysis code will be available upon reasonable request

## ACKNOWLEDGEMENTS

We are grateful to all members of the Nerurkar lab for insightful discussions and feedback. We are additionally grateful to Lianna Gangi and Clark Hung for assistance with biochemical quantification of collagen via OHP assay.

