## Supplemental Figure for "Elastic fibers define embryonic tissue stiffness to enable buckling morphogenesis of the small intestine"

**-**

**SUPPLEMENTAL DATA**

a.

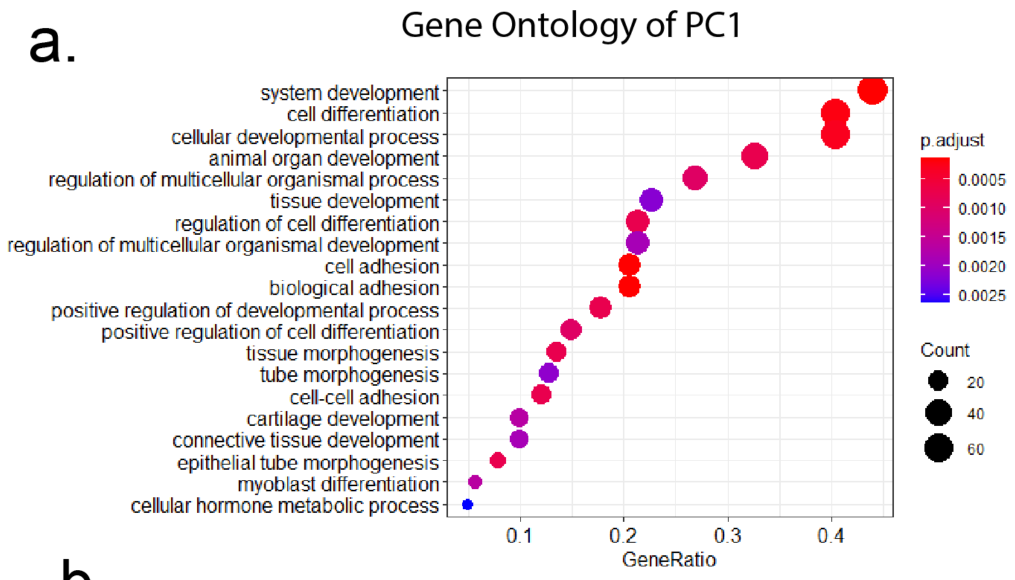


b.

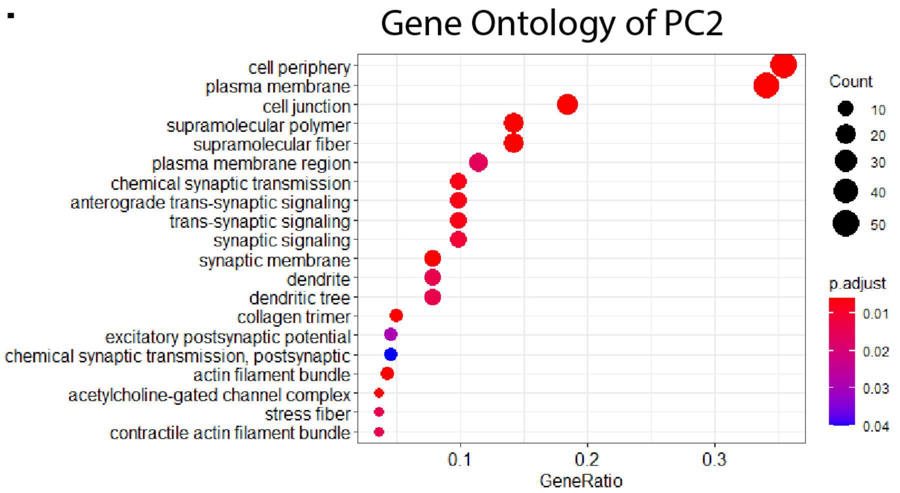

***Figure S1 (supplemental data) - bulk RNAseq***

**a.** Dot plot representing results of Gene Ontology analysis of 500 top genes driving Principal Component 1 of PCA **b.** Gene Ontology analysis of 500 top genes driving Principal Component 2, where ontologies related to nervous cells are enriched.

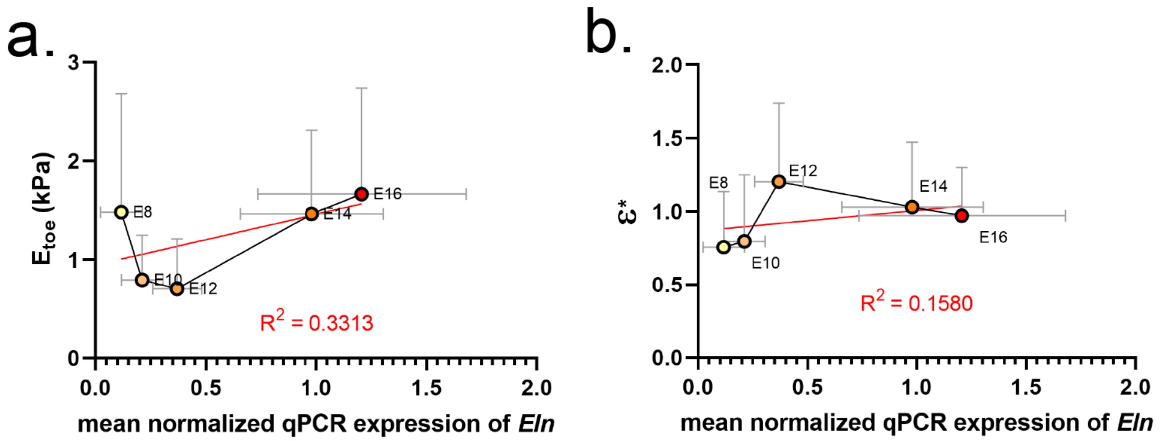


***Figure S2 (Supplemental data) - mechanical testing and Eln qPCR expression***

**a.** Weak/no correlation between expression levels of *Eln* and toe modulus ($E_{toe}$) in the mesentery between E8 and E16 ($R^2$ =0.3313) **b.** Weak/no correlation between expression levels of *Eln* and transition strain (ε*) in the mesentery between E8 and E16 ($R^2$ =0.1580).

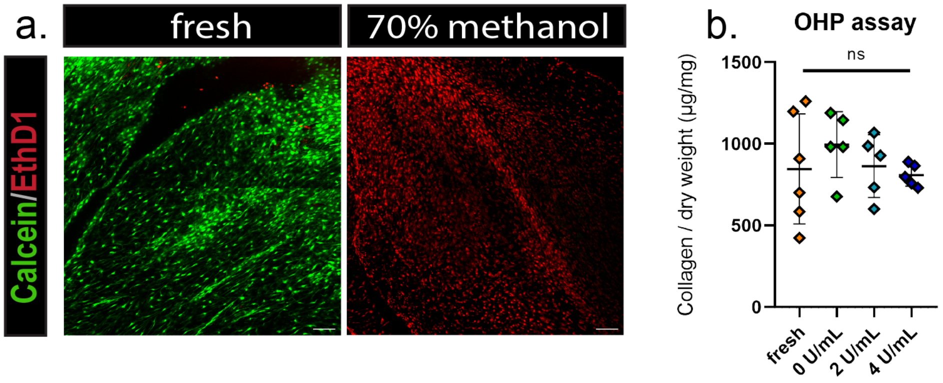


***Figure S3 - Elastase treatment controls***

**a.** Validation experiments for cell viability assay conducted on freshly dissected, untreated tissue (left) and tissue incubated with 70% methanol (right). **b.** Quantification of collagen content in control and elastase-treated mesentery samples evaluated by ortho-hydroxyproline (OHP) assay; n.s. = not significant.

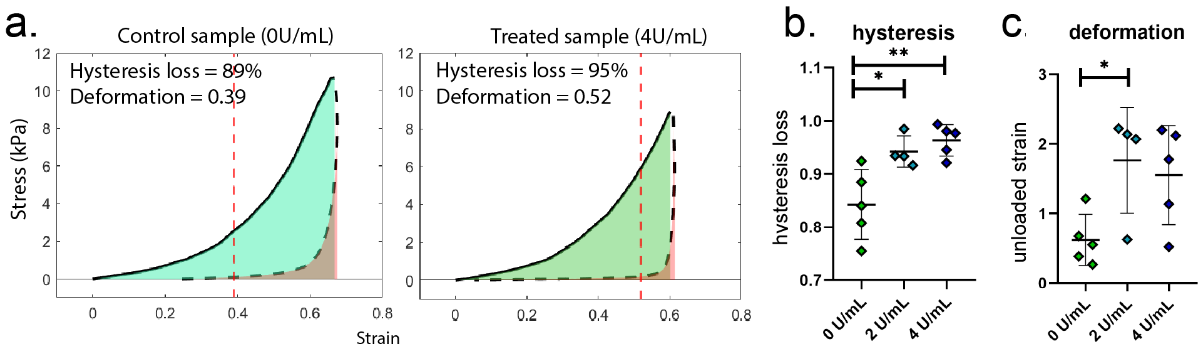


***Figure S4 - Elastase treatment reduces elastic energy storage in the mesentery.***

**a.** Example curves for loading (solid black line) and unloading (dashed black line) mechanical testing **b.** Hysteresis loss as fraction of initial hysteresis in elastase treated and untreated samples **c.** Deformation (remaining strain after unloading). n.s. = not significant ($p>0.05$); *p $<0.05$; **p $<0.005$.

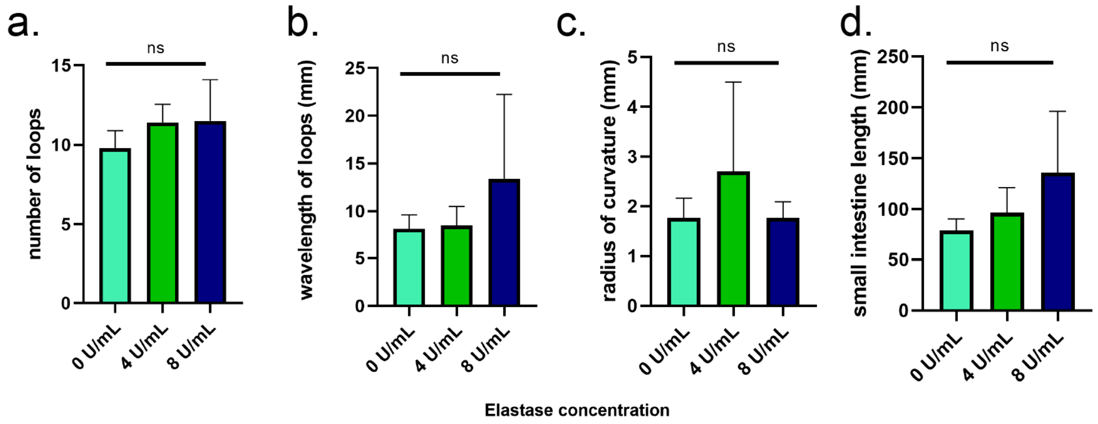


***Figure S5 - Elastase treatment of whole intestines does not significantly alter morphometry of loops***

**a.** Numbers of intestinal loops for control and elastase-treated intestines. **b.** Wavelength (intestinal length divided by number of loops). **c.** Mean radius of curvature of loops. **d.** Small intestine length. *n.s. = not significant (p>0.05).*
